# Statistical relationships across epigenomes using large-scale hierarchical clustering

**DOI:** 10.1101/2024.10.21.619460

**Authors:** Anastasiia Kim, Nicholas Lubbers, Christina R. Steadman, Karissa Y. Sanbonmatsu

## Abstract

Environmental toxins and pathogens can influence epigenetic modifications on chromosomes, thereby leaving trace evidence of exposures. However, the avalanche of epigenomic data is difficult to parse for biological interpretation given non-linear complex patterns and relationships. This attractive challenge in epigenomic data lends itself to machine learning for discerning infectivity and susceptibility. In this study, we explore over 3,000 epigenomes of uninfected individuals and provide a comprehensive characterization of the relationships among epigenetic modifiers, their modifiers, and specific immune cell types across all chromosomes using hierarchical clustering.

## I. Introduction

Epigenetic modification of the genome contributes to the function and physiology of all organisms. These covalent additions of functional groups can occur directly on DNA and histone proteins via chromatin modifying enzymes and are termed DNA methylation and histone (post translational) modifications, respectively. Epigenetic modifications influence various cellular and organismal processes ranging from basic biochemistry to immune function and even human behavior [Keverne et al., 2015, Obata et al., 2015, Tiffon, 2018]. The booming growth of the epigenomics field—that is, the study of epigenetic modifications across the entire genome—can be attributed not only to growing interest of the scientific community but also the recent onslaught of advances in sequencing technology [Callinan and Feinberg, 2006, Clark et al., 2016]. As sequencing platforms continue to evolve, the number of epigenomic datasets continues to grow, and such data is housed within the ENCODE, SRA, and GEO databases [Barrett et al., 2005, Leinonen et al., 2010, Consortium et al., 2012]. These datasets provide a wealth of information, particularly for understanding the contribution of aberrant epigenetic modifications in disease states, such as cancer, and the impact of environmental exposures on function and behavior [Wolffe, 2001, Jirtle and Skinner, 2007, Hou et al., 2012, Ilango et al., 2020, Perera et al., 2020]. However, the complexity and sheer enormity of these datasets precludes definitive biological interpretation; as such, this is a unique opportunity to utilize machine learning approaches to identify patterns and signatures that may reveal mechanisms of normative and/or aberrant physiological processes. In particular, the immune system utilizes epigenetic modifications and chromatin remodeling processes to respond to threats: the orchestrated epigenomic regulation of immunity, both innate and adaptive function, presents a large complex story to unravel. The details and precise mechanisms of this regulation would provide useful information for prediction, early detection, and development of effective treatments [Fernández-Morera et al., 2010, Janson and Winqvist, 2011].

The machine learning and data science revolution of the past decade has now been directed at a great many scientific endeavors. Unsupervised learning methods have been used extensively in genomics and epigenomics studies, with particular focus on DNA methylation. In particular, Titus et. al showed that variational autoencoders can learn latent DNA methylome (methylation across the genome) representation that can be used for lower dimensional epigenetic analyses [Titus et al., 2018]. Zamanighomi et. al [Zamanighomi et al., 2018] identified clusters of informative peaks for single-cell methylation data using unsupervised clustering. Hierarchical clustering was employed in studies to analyze DNA methylation patterns: Virmani et. al [Virmani et al., 2002] showed that DNA methylation patterns differ between lung cancer cell lines, while Lin et. al revealed tumor-specific hypermethylated clusters and expressed breast cancer genes [Lin et al., 2015]. Hierarchical clustering of diffuse large B-cell lymphoma based on the extent of DNA methylation variability identified novel epigenetic clusters [Chambwe et al., 2014]. These important studies demonstrate the utility of hierarchical clustering and unsupervised learning on identifying patterns based on DNA methylation. Yet, few studies have addressed histone modifications and associated chromatin regulatory proteins. Compared to the simplicity of DNA methylation datasets, the sheer number of histone modifications, variations, and interactions creates a complex pattern recognition challenge [Seligson et al., 2005, Xi et al., 2018]. Thus, in this work, we turn to the tools of data science to meet this challenge, focusing on using hierarchical clustering of large scale epigenomic data, to provide a birds-eye-view of the relationships between histone modifications and associated chromatin regulatory proteins across a variety of immune cell types. We collated a dataset comprised of 3111 whole-genome histone and chromatin modifiers samples totaling approximately 1.5 TB of data from the Encode Consortium [Consortium et al., 2012]. We note that this paper does not delve into a detailed analysis for each chromosome; rather, we present a methodological framework which serves as a foundation for future, more detailed investigations of individual chromosome analysis.

## II. Data description

To understand how epigenetic modifications of chromosomes are impacted by pathogen exposures resulting in activated immune function, it is necessary to first establish a baseline from uninfected individuals to understand the relationships between epigenetic modifications and associated chromatin regulatory proteins. We acquired 3111 ChIP-seq samples on various histone modifications and chromatin binding proteins from cells derived from healthy, normal subjects (uninfected) from the Encode project [Consortium et al., 2012]. These samples were based on either the hg19 (1479 samples) or the hg38 (1632 samples) human genome reference assembly, with each sample characterized by a specific epigenetic modifier in a specific immune cell type. In addition to being classified as either a histone or not, each epigenetic modifier was further characterized by the factor it is associated with and its activity. Factor represents different roles and mechanisms in the epigenetic regulation process and the activity of each modifier indicates its role in regulating gene expression. In total, there were 28 unique cell types and 28 unique epigenetic modifiers in the h19 (GRCh37) assembly samples and 14 unique cell types and 68 unique epigenetic modifiers in the h38 (GRCh38) assembly samples. Each sample was stored in a BigWig file, which contained the signal p-value, which is -log10 (p-value). The signal p-value tests the null hypothesis that the signal at a particular genomic loci is present in the control. BigWig files are a format used to store large sets of genomic data that would otherwise take up too much space and be cumbersome to process. They allow for efficient retrieval of subsets of data, making it easier to access and visualize specific regions of a genome without the need to process the entire file.

## III. Results

In this section, we present the results of our analysis, which primarily focused on the hg38 (GRCh38) genome assembly reference datasets and chromosome 6, known for its abundance of immune-related genes [Shiina et al., 2009]. Additionally, we have validated our findings on samples aligned to the hg19 (GRCh37) genome assembly to ensure the robustness of our methodology (Fig. 1).

**Fig. 1.**
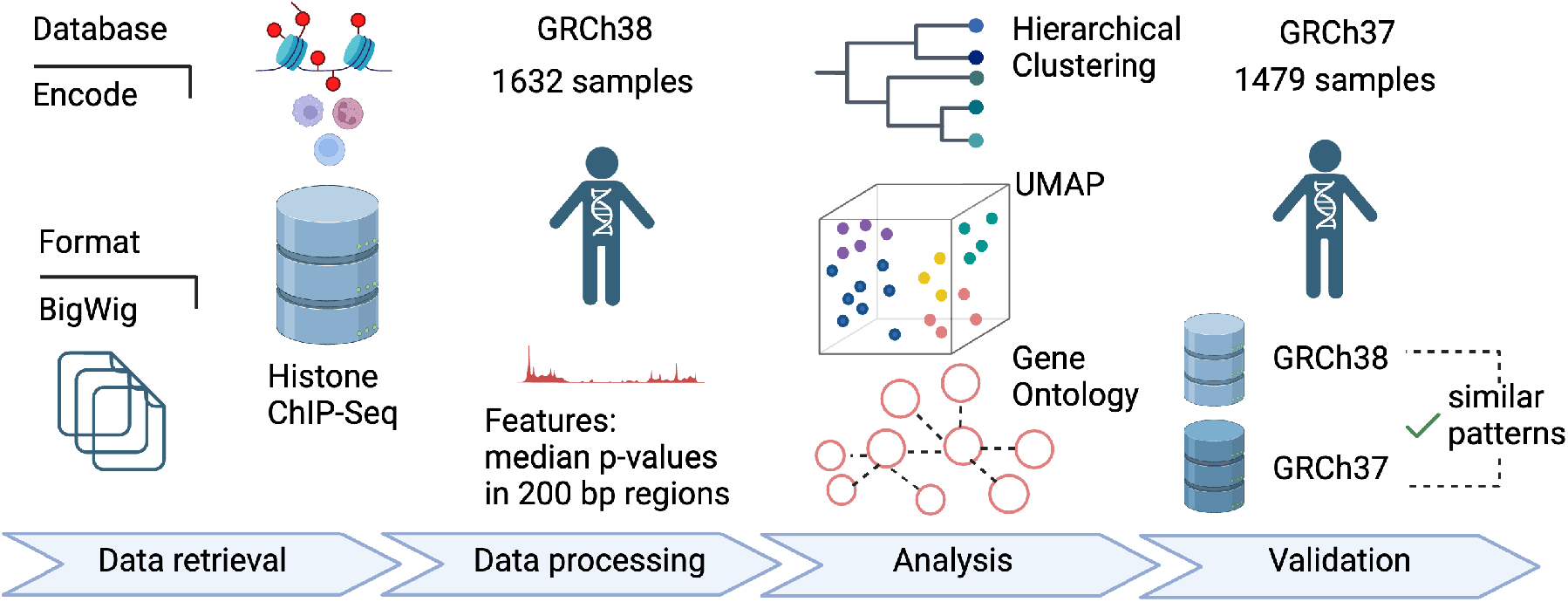
A flowchart of our approach. Note that datasets from the two genome reference assemblies are different samples.

### A. Numerous known genes, identified among important features, drove the clustering

We observed the abundance of important known genes in most of the clusters, indicated by the color-coded branches in the hierarchical clustering dendrogram of chromosome 6. The pie charts in the dendrogram, each varying in the number of colors, show the count of distinct epigenetic modifiers within each cluster (Fig. 2). Several pie charts consisted of only one or two colors, signifying that clusters containing at least 30 important known genes, were characterized by one or two specific epigenetic modifiers, respectively. This pattern suggested that a considerable number of epigenetic modifiers tended to group together, reflecting a clustering of samples influenced predominantly by the specific traits of these modifiers.

**Fig. 2.**
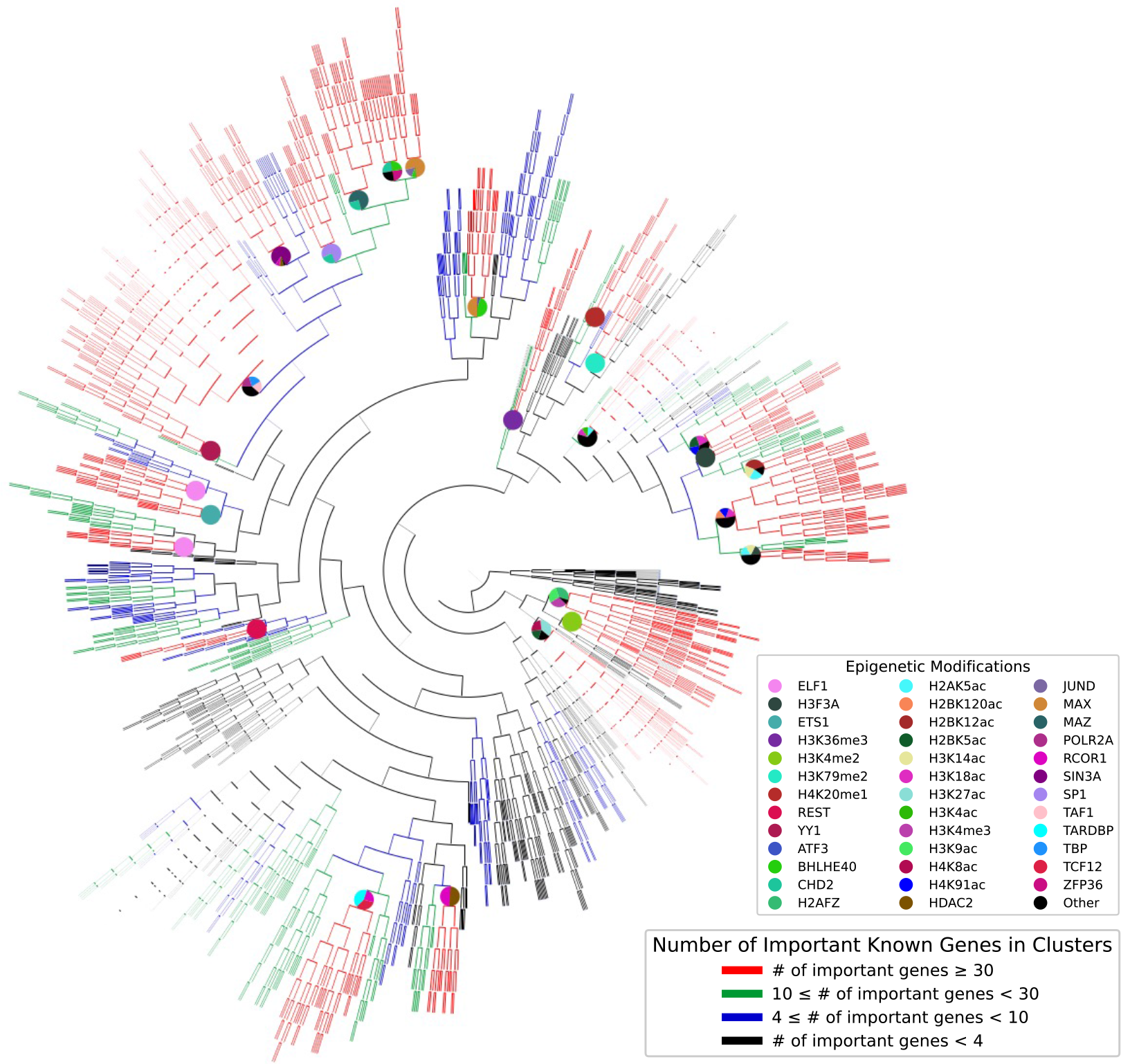
Dendrogram of representative chromosome (chromosome 6). Here, red branches denote clusters where the number of important (common) known genes in all samples are at least 30. The labels of each of such clusters are also represented by the pie charts, where the number of colors indicate how many distinct epigenetic modifiers are in the cluster. Note if there are more than 4 distinct modifiers, the least frequent ones are combined in one pie chart slice.

### B. Samples showed a tendency to cluster together according to the epigenetic modifiers first and then by the cell types

While the dendrogram hinted that similar epigenetic modifiers tend to cluster together, UMAP plots not only confirmed this tendency but also revealed that samples clustered first by the modifiers and then by the cell types. However, some samples characterized by the certain cell types like peripheral blood mononuclear cells, common myeloid progenitors, CD34-positive cells, neutrophils, and CD14-positive monocytes showed a tendency to cluster primarily by these cell types (Fig. 3). In contrast, for other cell types, the clustering was predominantly driven by modifiers (Fig. 4). The clustering patterns observed in UMAP plots suggested samples tended to group together based on the most distinctive features in the dataset. In this context, if a specific cell type has more unique and defining characteristics than an epigenetic modifier, we expect the samples will cluster according to cell type and vice versa. These several cell types (peripheral blood mononuclear cells, common myeloid progenitors, CD34-positive cells, neutrophils, and CD14-positive monocytes) that tended to form clusters were the most prevalent within the dataset which might had an effect on the UMAP plots. In the hg38 genome dataset, the epigenetic modifiers were evenly distributed, with each of the 68 types represented by 24 samples. However, the representation of cell types in the dataset showed a different pattern. Two out of the 14 cell types—common myeloid progenitor and CD34-positive peripheral blood mononuclear cells—accounted for about a third of the dataset, indicating a skewed distribution in the representation of cell types.

**Fig. 3.**
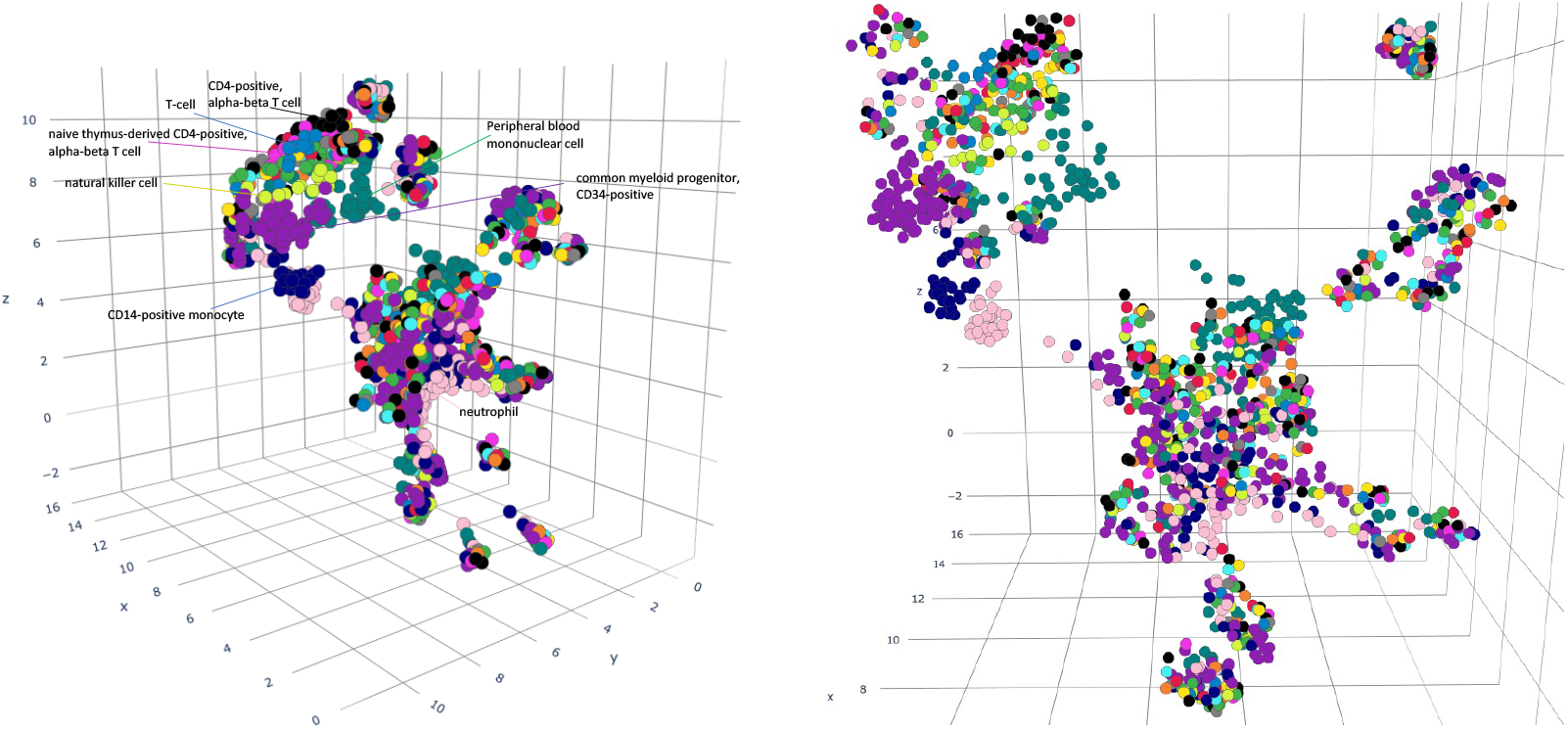
UMAP projections from different angles of chromosome 6. Each sample is color-coded to indicate its associated cell type. Each color represents one of the 14 cell types. The most visually distinct samples are for clarity. The axes represent the components or dimensions that UMAP has reduced the data to.

**Fig. 4.**
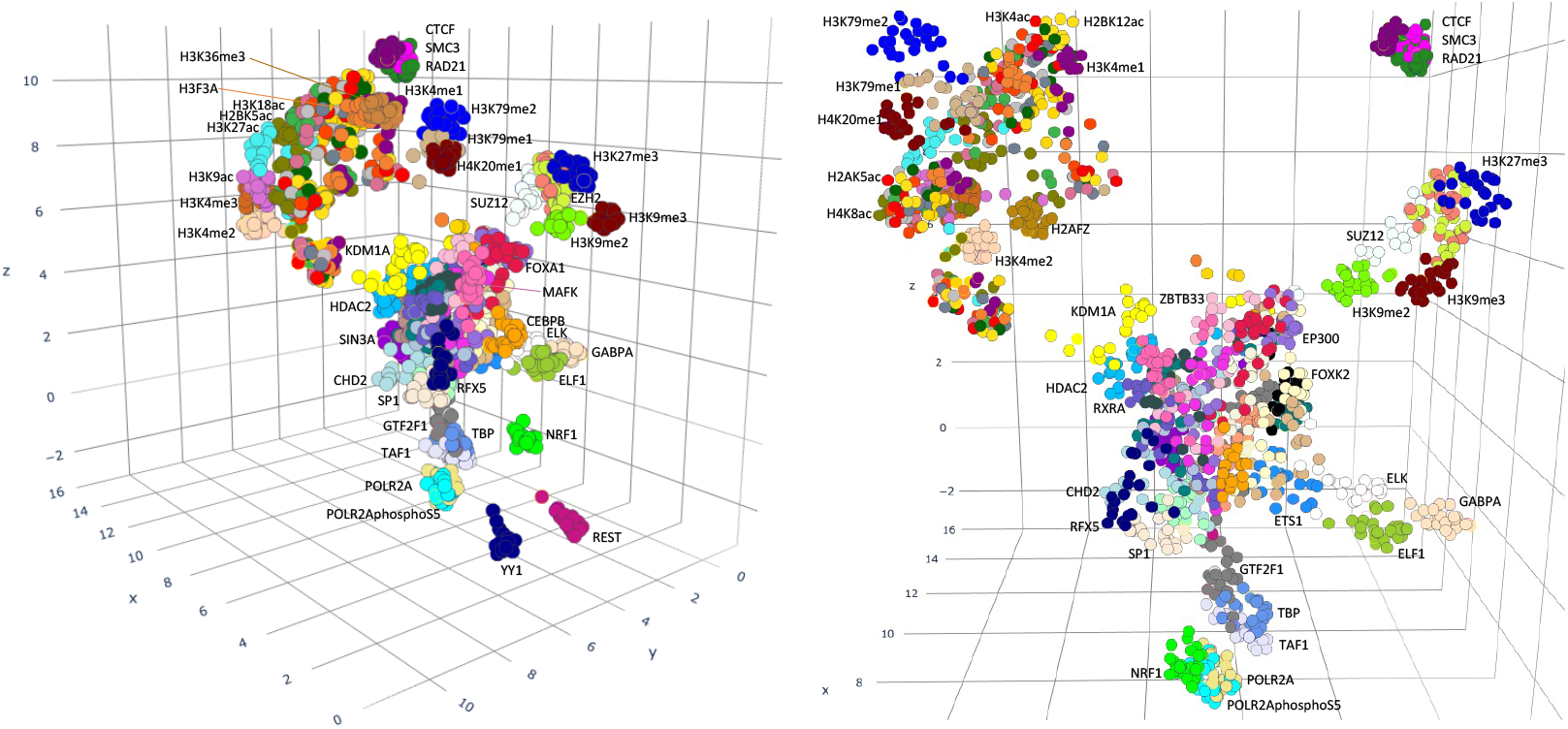
UMAP projections from different angles of chromosome 6. Each sample is color-coded to indicate its associated epigenetic modifier. Given that there are 68 unique modifiers in the dataset, some colors may not be distinctly differentiable to the eye. The most visually distinct samples are labeled for clarity. The axes represent the components or dimensions that UMAP has reduced the data to.

The main distinction in clustering was notably between histone and non-histone modifiers (Fig. 5). Modifiers typically clustered based on their activity (such as permissive or repressive) and factor (transcription factor, chromatin modifier, histone modification, etc.). However, a more pronounced distinction was observed between histone permissive and histone repressive modifiers compared to that between non-histone permissive and non-histone repressive modifiers (Fig. 5).

**Fig. 5.**
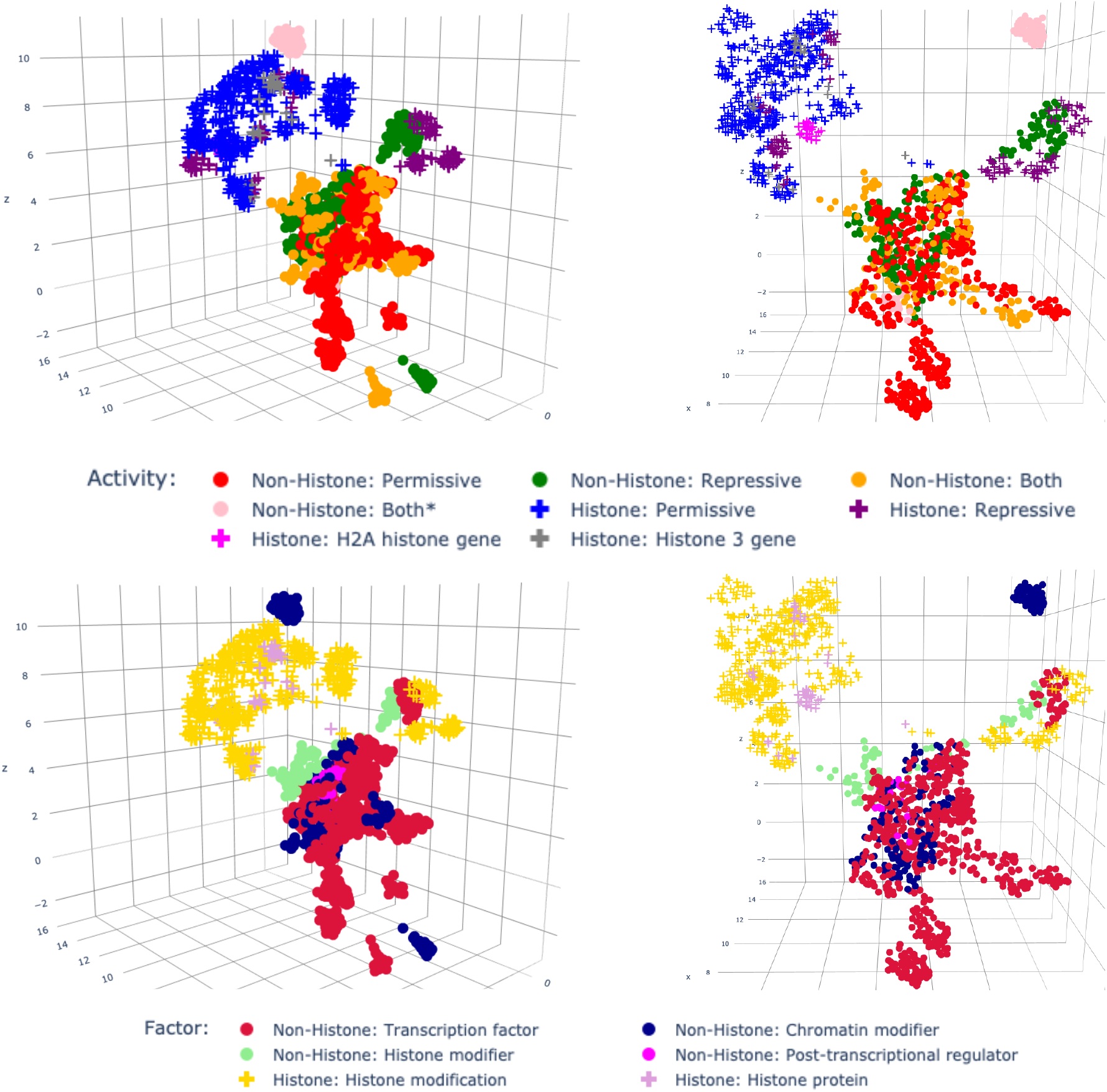
UMAP projections from different angles of chromosome 6. Each sample is color-coded to indicate its associated factor and activity. The shapes of the markers indicate whether each epigenetic modifier is a histone or not. The axes represent the components or dimensions that UMAP has reduced the data to.

### C. Clustering behavior was consistent across all chromosomes

We were interested in identifying which features drove clustering and whether they were linked to specific genes. To achieve this, we needed to examine each cluster that can be obtained by cutting the dendrogram at choosen height. Our observation of a roughly constant branching fraction across the dendrograms suggested a similar structure across all chromosomes (Fig. 6).

**Fig. 6.**
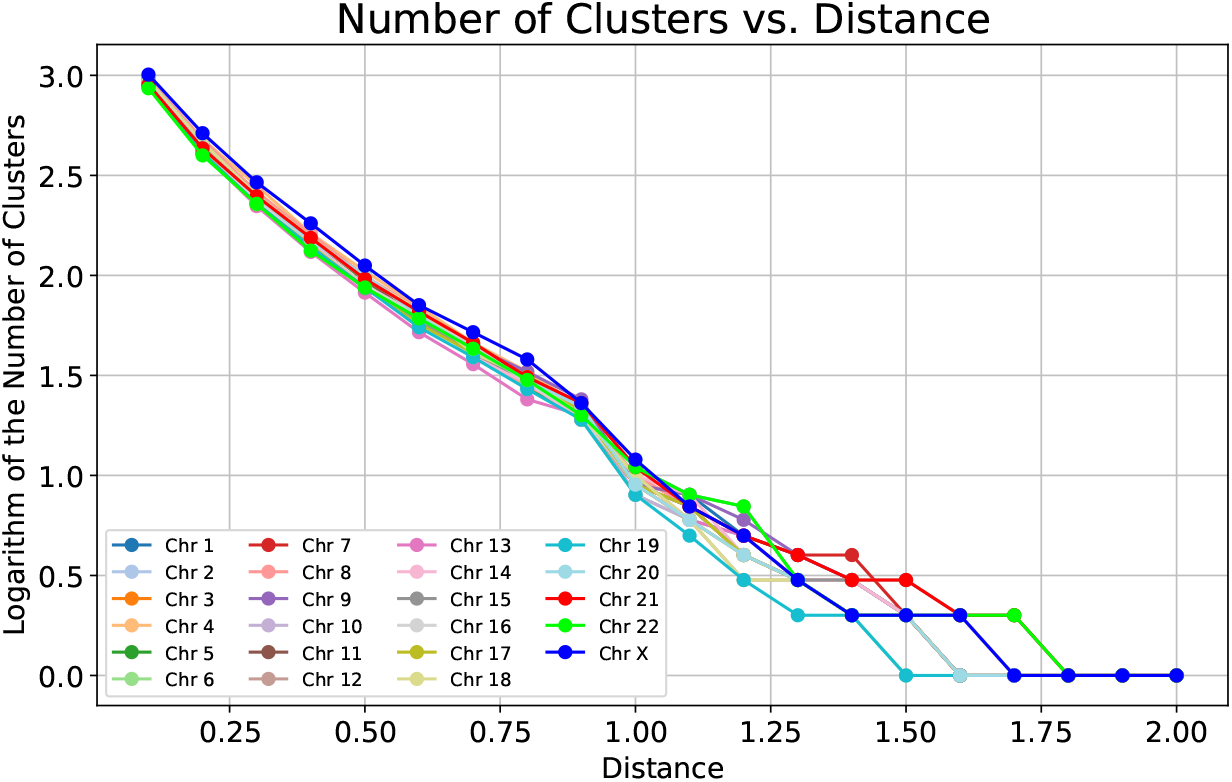
The number of clusters (*log*_10_) found using the hierarchical clustering on each chromosome, as a function of the distance (1 - Pearson correlation) used to cut the hierarchical clustering. All chromosomes have a similar structure with a roughly exponential decay in clusters. This corresponds to a roughly constant branching fraction for the dendrogram.

Additionally, plotting entropy against distance offered further insights, especially when comparing different chromosomes (Fig. 7). We observed how entropy varied as a function of the cut point; where entropy decreased rapidly, the dendrogram cuts were effectively dividing clusters into more similar labels. The entropy behavior was analyzed with respect to the epigenetic modifiers, activity and factor associated with each epigenetic modifier, and cell types (Fig. 7). On the right-hand limit, each dendrogram corresponded to one unique cluster which contained all of the samples, and thus all chromosomes had exactly the same entropy value, which was the entropy of the dataset. On the left-hand limit, each cluster had exactly one sample, and so the entropy was precisely zero for each cluster, and thus zero for each dendrogram. In between, the variation in entropy as a function of the cut point was observed. Cross-referencing with the number of clusters plot (Fig. 6), we observed that the entropy started to diminish at an early stage, marked by a higher distance values, when there were a few clusters. It made a more dramatic jump downwards around 10 clusters (where the distance was around 1) for the activity, factor, and epigenetic modifier plots. For the cell type, the entropy curves were qualitatively different from the entropy curves associated with the epigenetic modifiers. These curves were approximately flat from the cut point 1 upwards, indicating that the highest levels of the dendrogram did not make any noticeable effect on the cell line distribution. By comparing the behavior of black curve corresponding to the randomly shuffled labels to the original entropy curves, we observed that there was less evident structure in the cell type behavior meaning that clustering by the cell types was less meaningful than the clustering by other identifiers. We also used the Adjusted Rand Index (ARI) to compare dendrograms across all chromosomes. The ARI, notable for its independence from specific labels, unlike entropy, facilitates a more comprehensive comparison of clustering patterns. Our analysis revealed that the pairwise ARI values in chromosome comparisons ranged from approximately 0.5 to 0.9. This range suggested that clustering patterns were more similar than dissimilar between all chromosomes.

**Fig. 7.**
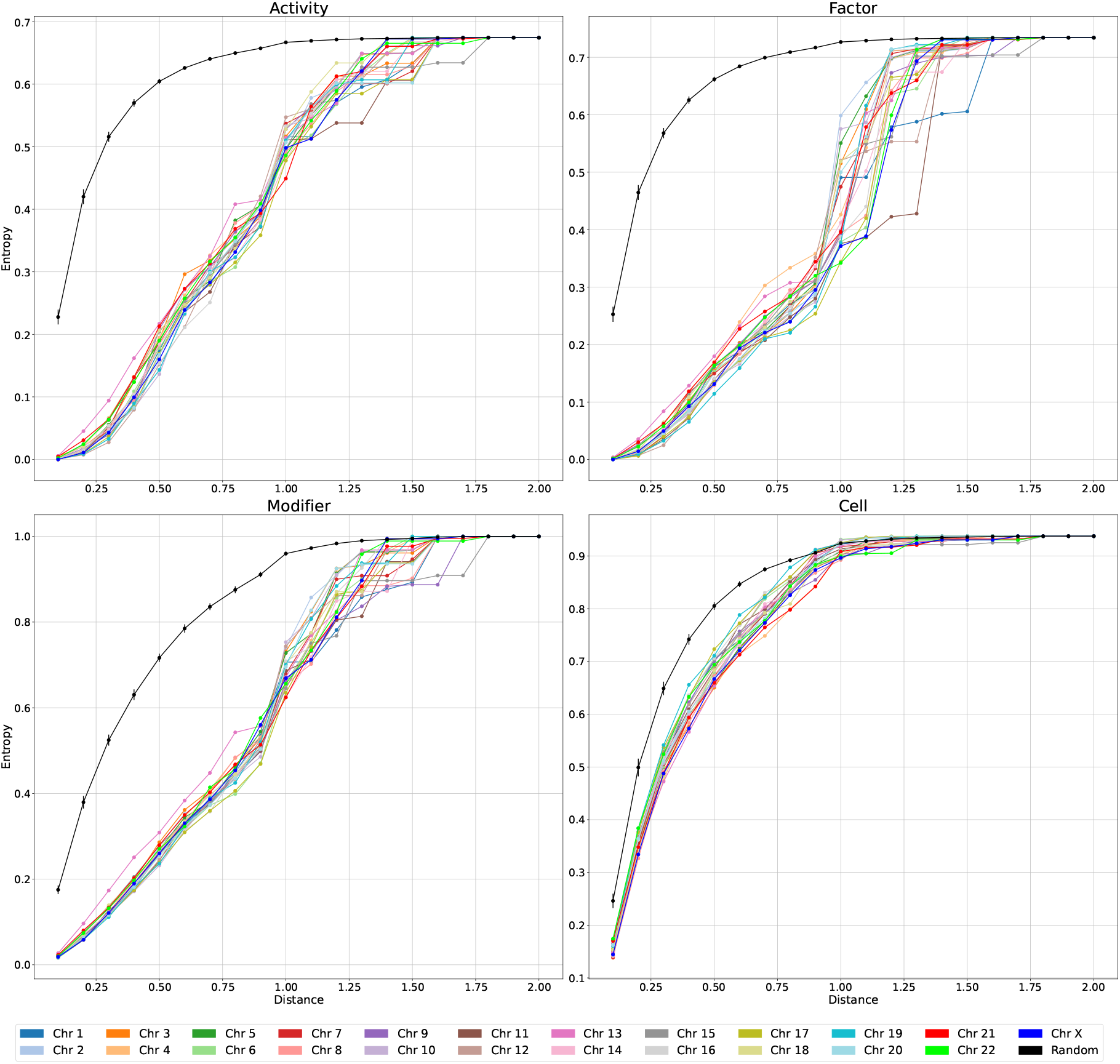
Weighted entropy vs. Distance for true and random labels. The entropy values were calculated with respect to activity (permissive/repressive/both), type of factor (transcription/histone modifier/chromatin modifier/others), modifier (68 types), and cell type (14 types) for each chromosome, as a function of the distance (1 - Pearson correlation) used to cut the hierarchical clustering dendrogram. Black lines on the plot correspond to the entropy values when all labels in the dendrogram were randomly shuffled. Calculations were performed 100 times. Note that the y-axis is scaled differently in each plot.

### D. Several modifiers co-occured together in the same clusters across all chromosomes

The co-occurrence matrix heatmap revealed how epigenetic modifiers were related, specifically whether they tended to be present together in the same clusters across all chromosomes (Fig. 8). The co-occurrence patterns of modifiers showed that Histone 2 and 3 acetylation modifiers commonly co-occurred. The trio of RAD21, SMC3, and CTCF was also frequently found to co-occur, consistent with their participation with the cohesin complex. POLR2A and its phosphorylated form, POLR2AphosphoS5, often clustered together with the pair of TAF1 and TBP. Also, TAF1 and TBP were frequently found in the same cluster with GTF2F1 consistent with their involvement in RNA Polymerase II (Pol II) transcription. EZH2, alongside its phosphorylated variant, EZH2phosphoT487, co-occurred with SUZ12 and H3K27me3 consistent with the role of Polycomb Repressive Complex 2 (PRC2) complex depositing the H3K27me3 modification. Another set of modifiers that tended to appear together were BHLHE40, MAX, CHD2, and MAZ; BHLHE40 also co-occured with the pair of USF1 and USF2. Another notable co-occurence included TCF12, RCORI, ZFP36, RXRA, and HDAC2. Also, EP300 and FOXA1 showed a tendency to cluster within the same cluster. Interestingly, these clustering trends were not confined to chromosome 6 alone; many of the modifiers that co-occurred on chromosome 6 were also found clustering together on other chromosomes. This pattern was evident in both UMAP plots and co-occurrence heatmaps, indicating a broader chromosomal consistency in the clustering behavior of these epigenetic modifiers (Fig. 4, 8).

**Fig. 8.**
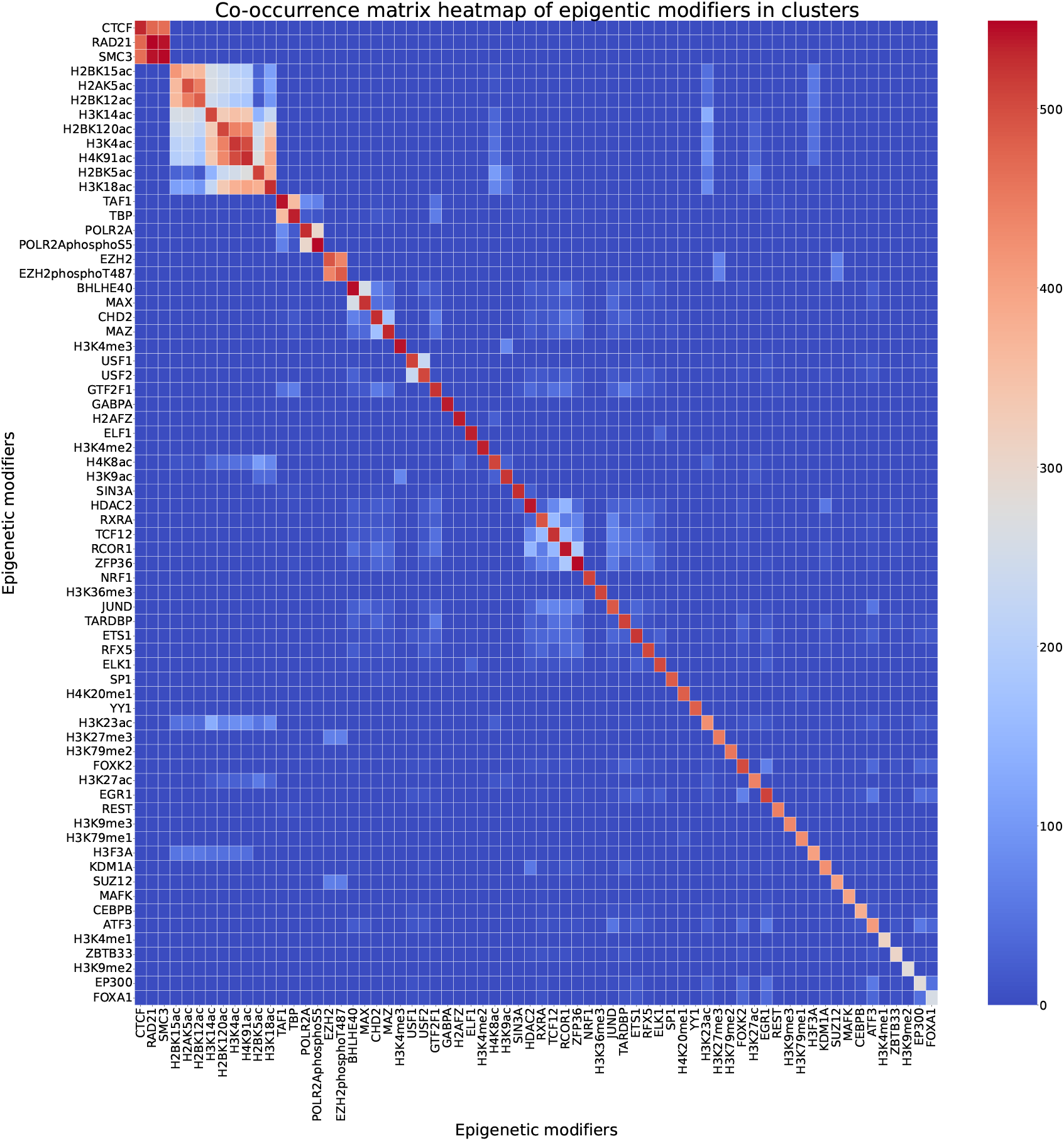
Co-occurrences of epigenetic modifiers within the clusters across all chromosomes. The intensity of red coloration corresponds to higher frequencies (sums of counts) of co-occurrences within the same clusters, following a dendrogram cut-off at a correlation distance below 0.3. Only clusters with a minimum size of 4 samples were chosen. The heatmap highlights the top 50 co-occurring modifiers.

### E. Many of the known genes were associated with microRNAs and snoRNAs

Finally, we connected important genes found across all chromosomes to the Gene Ontology and the KEGG pathway databases. We found several significant GO terms that were enriched in our important genes (Fig. 9), including those involved with chromatin remodeling and immune responses. For example, the GO term “Innate immune response to mucosa” was aligned to the h2BC10, H2BC11, H2BC6, H2BC7, H2BC8, and RNASE3 genes, which are directly related to the immune system processes. Meanwhile, other terms were related to fundamental cellular processes and structures involved in various aspects of immune cell function. KEGG pathway analysis revealed important genes that were connected to microRNAs, which were best annotated and studied in the context of cancer [Virmani et al., 2002], and therefore identified as the most significant in our analysis. We also identified other microRNAs related to roles of the spliceosome, immune function (i.e., neutrophil extracellular trap formation and systemic lupus erythematosus), and alcoholism (Fig. 10).

**Fig. 9.**
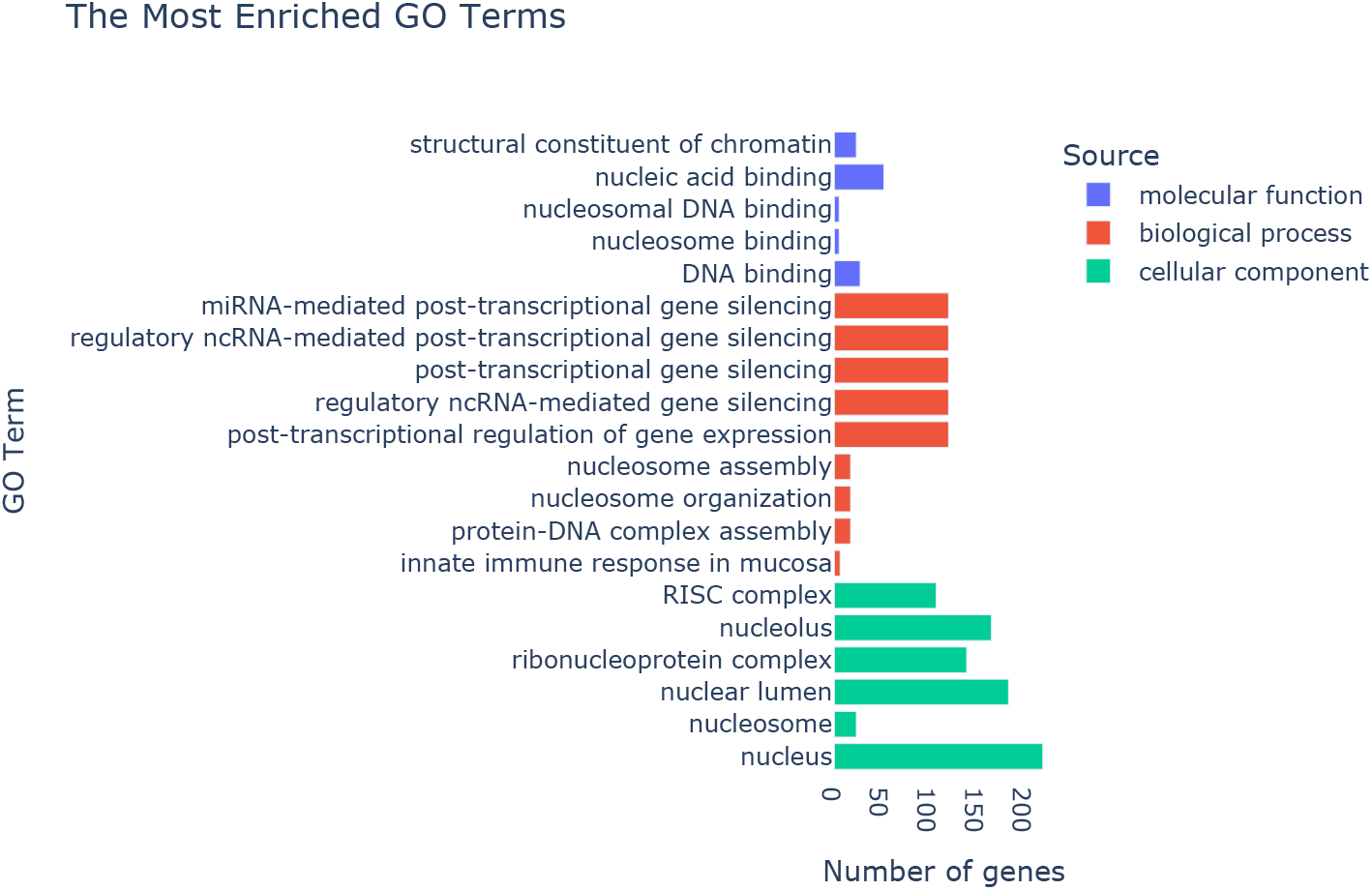
Gene ontology results for the important genes found across all chromosomes (using gProfiler tool). Several molecular functions, biological processes, and cellular components are identified. Innate immune response in mucosa, the body’s early defense mechanism in mucosal tissues, is associated with genes such as H2BC10, H2BC11, H2BC6, H2BC7, H2BC8, and RNASE3 genes.

**Fig. 10.**
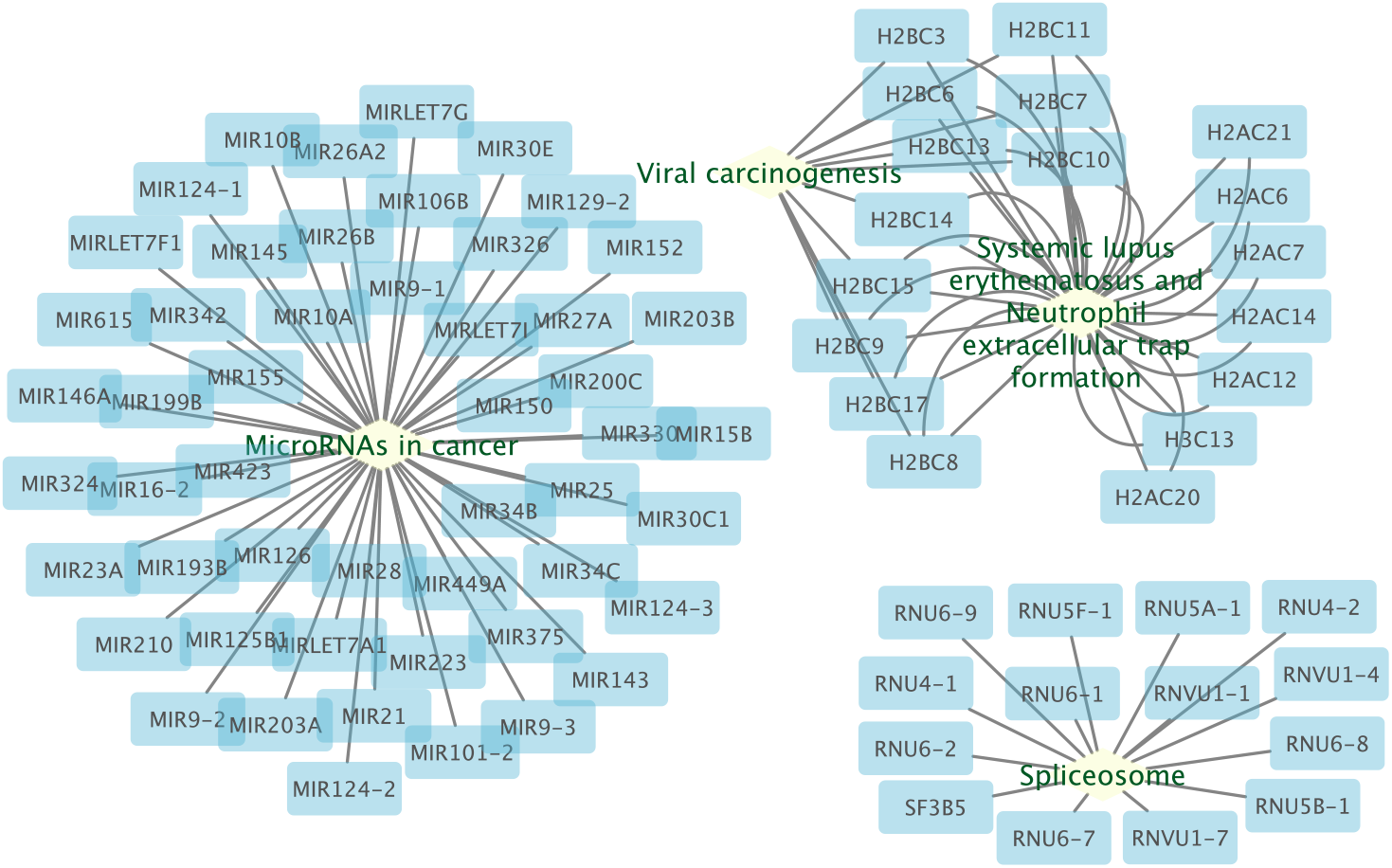
KEGG pathway network of important genes (*p <* 0.05 in all samples in cluster) gathered across all chromosomes. The functionality grouped network is visualized using Cytoscape based on the connectivity between pathways and genes. Many of the 51 microRNAs genes are annotated as involved in cancer pathways, as those are the most well studied. Some small nuclear RNAs are connected to the spliceosome. Many genes in the histone H2B family are found to be dysregulated in viral-induced cancers. These H2B genes and several others from the H2A histone family are linked to neutrophil extracellular trap formation, as histones play a major role in the NETs (Neutrophil Extracellular Traps) framework to halt invaders. Dysregulation of such processes occurs in autoimmune diseases, such as systemic lupus erythematosus, and in cancer, diabetes, and alcoholism (not shown in the plot).

### F. Validation on another dataset revealed similar patterns

We evaluated our approach on another dataset consisting of 1479 samples aligned to the hg19 genome assembly. To make comparison fair, we randomly selected 600 samples from each hg19-aligned and hg-38-aligned dataset, resulting in an equal number of 14 cell types and 28 modifiers. We observed similar branching factors on the entropy plot for all type of labels (Fig. 11). However, all entropies were lower for the hg19-aligned dataset than for the hg38-aligned dataset, making it hard to tell whether the observed differences were due to the datasets themselves or if the difference between hg19 and hg38 genome assemblies played a significant role. The co-occurrence matrix heatmaps revealed that the same epigenetic modifiers tended to cluster together regardless of the dataset, with the exception of H4K8ac, which co-occurred with a bunch of different modifiers in the hg19-aligned dataset but not in the hg38-aligned one (Fig. 12).

**Fig. 11.**
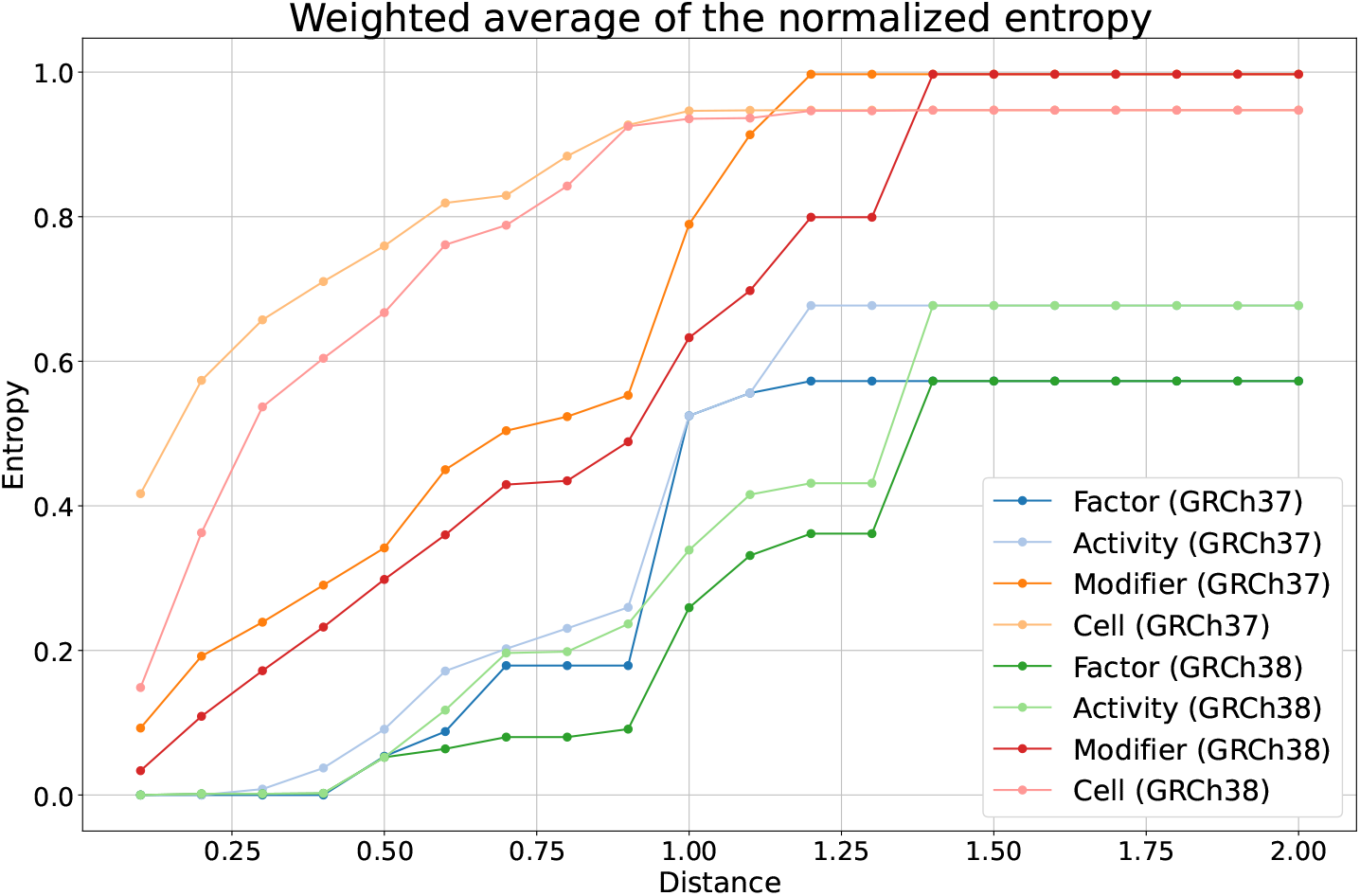
Weighted entropy vs Distance. The calculated entropy values for each chromosome for hg19 and hg38 datasets, with respect to activity type, factor, modifier, and cell type, as a function of the distance used to cut the hierarchical clustering dendrogram. To ensure a fair comparison, we randomly selected 600 samples from each dataset, resulting in an equal number of 14 cell types and 28 modifiers.

**Fig. 12.**
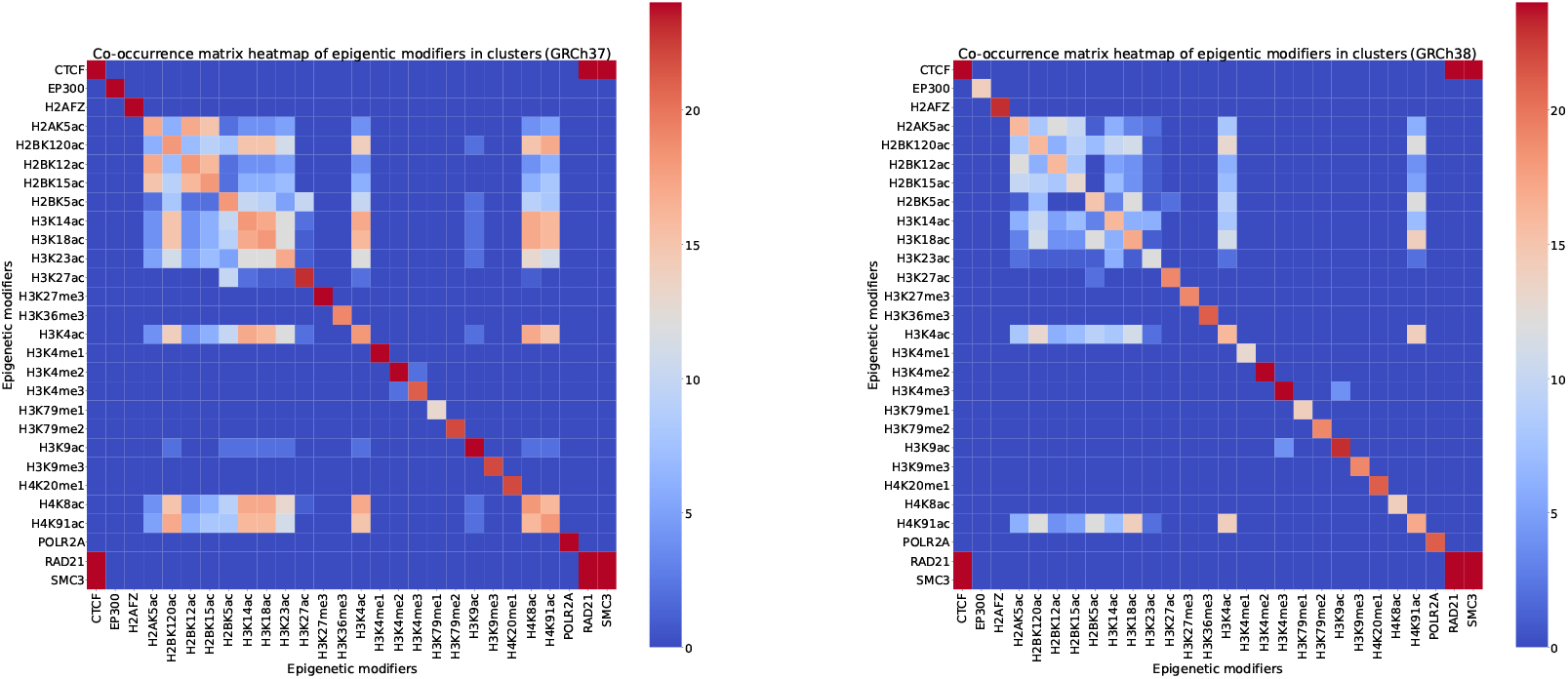
Heatmaps that illustrate the patterns of epigenetic modifier co-occurrences within the clusters in chromosome 6 derived from distinct datasets of the human genome assemblies hg19 (left) and hg38 (right) datasets. The intensity of red coloration corresponds to higher frequencies (sums of counts) of co-occurrences within the same clusters, following a dendrogram cut-off at a correlation distance below 0.3. Only clusters with a minimum size of 4 samples were chosen. The heatmaps highlight the top 50 co-occurring modifiers. The heatmaps exhibit a similar pattern of co-occurrences, indicating that certain epigenetic modifiers tend to co-occur together regardless of the dataset. To ensure a fair comparison, we randomly selected 600 samples from each dataset, resulting in 28 unique modifiers.

## IV. Methods

Our analysis focused primarily on samples aligned to the hg38 (GRCh38) genome assembly reference. To evaluate the generalizability of our methodology, we also applied our analysis to samples aligned to the hg19 (GRCh37) genome assembly. This step was taken to confirm that the patterns we observed were not unique to a specific dataset. For each chromosome, we divided the genome into 200 bp regions and extracted the median signal p-value for each region. These median values were then transformed into regular p-values. Using these 200 bp regions as features, we conducted hierarchical clustering on the samples.

### A. Hierarchical Clustering

Hierarchical clustering has gained popularity due to its straightforward implementation and visualization capabilities [Murtagh and Contreras, 2012]. The results of agglomerative clustering might differ based on the choice of distance metrics and linkage techniques, which dictate cluster merging decisions. We conducted hierarchical clustering using correlation distance (1 *− r*, where *r* is the Pearson correlation coefficient) and complete linkage clustering. While correlation distance is not strictly a distance in mathematical terms, it allows to derive clusters that focus on relationships between features (regions of the genome in our case) rather than absolute magnitudes. In our study, this distance was particularly useful for capturing trends where there were peaks in regions of the genome across multiple samples. We used complete linkage, where the distance between clusters is the largest distance between any two members of the cluster, as it is robust to outliers and tends to produce tighter clusters, aligning with our objective of clustering samples based on similar feature (regions of the genome) behaviors.

As hierarchical clustering is an unsupervised learning technique, it naturally leads one to the question: how can we assess the quality of the clusters especially when too many clusters were formed? The optimal clustering scenario occurs when the within-cluster distance is minimized while the between-cluster distance is maximized. Typically, cluster validation employs internal or external metrics [Murtagh and Contreras, 2017]. Given that we used correlation rather than, for example, Euclidean distance, many traditional internal validation metrics become unsuitable, as they often rely on minimizing the sum of squared errors. The challenge with correlation is that one can not merely average correlations in the same way distances are averaged. Furthermore, internal metrics can exhibit biases and might not always offer the best validation. On the other hand, external validation methods such as the adjusted rand index, Jaccard coefficient, entropy, purity, and Fowlkes-Mallows index require a known ground truth for comparison, which might not always be available, as in our case. However, a practical approach in our study involved leveraging the comparative analysis of different chromosomes and genome assemblies using entropy and adjusted rand index. For example, by treating one chromosome (or genome reference assembly) as a “ground truth”, we were able to compare the clusterings performed on two different chromosomes (or assemblies) with each other to determine their similarity.

### B. Entropy-based analysis across chromosomes

We used the entropy to compare the branching patterns of the dendrograms between chromosomes. The entropy for each cluster was defined as the weighted entropy 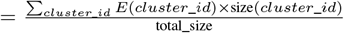 where 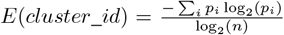 where *p*_*i*_ is the proportion of the *i*th label in the cluster and *n* is the number of unique labels. *E*(*cluster id*) is the entropy of a particular cluster, *size*(*cluster id*) is the number of elements in that cluster, and *total size* is the total number of elements across all clusters. In such way, we normalized the entropy to be between 0 and 1 which allowed for fair comparison across clusters with different numbers of labels, focusing on how evenly distributed the labels are, rather than just how many there are in a specific cluster. The weighted entropy over the clusters is defined so that each cluster is weighted by its size to ensure that the contribution of each cluster to the overall entropy is proportional to its representation in the data.

To validate our findings, we calculated the entropy for datasets with randomly shuffled labels, repeating the process 100 times to obtain an averaged result. This procedure was implemented to determine the presence of any underlying structure in the data to ensure that clustering did not happen by chance. In this study, entropy served as a quantitative measure of the level of variability within each cluster in the dendrogram, with lower entropy values indicating more homogeneous clusters. For each chromosome, we performed a detailed analysis by calculating entropy values at various cut points within the dendrogram. This was done separately for each case, depending on how the samples were labeled: as epigenetic modifiers, cell types, factors, or by their activity. By assessing entropy across these different labeling scenarios, we gained insights into how the clustering patterns varied depending on the specific characteristics assigned to the samples.

### C. Adjusted rand index analysis across chromosomes

In our study, alongside the entropy-based analysis where each sample was assigned a specific label type (modifier, cell type, factor, or activity), we also calculated the adjusted rand index (ARI) to compare the dendrograms across all chromosomes. The advantage of using ARI lies in its independence from specific labels, allowing for a broader comparison of clustering patterns across different chromosomes. The ARI is a corrected version of rand index, adjusted for chance, which accounts for the likelihood that two samples might end up in the same cluster purely by chance. This makes the ARI a more reliable measure in situations in which random agreement is more likely: cases where the dataset is large or the number of clusters is high. The ARI has a value close to 0 for random clustering and 1 for perfect clustering agreement.

### D. Investigating clusters: dendrogram and UMAP visualization

To understand the patterns within each chromosome of our dataset independently, we analyzed clusters in dendrograms and UMAP (Uniform Manifold Approximation and Projection) plots. Additionally, we examined the branching fraction in the dendrograms for each chromosome, where we analyzed the relationship between the number of clusters and the distance at which the dendrogram is cut. This assessment helped in understanding the clustering dynamics at different levels.

In addition to analyzing clustering patterns, we also sought to identify the most important (that is, common) regions in those clusters. We defined region of the genome as important in a certain cluster if its p-value was less than 0.05 in all samples within that cluster. For every 200 bp region marked as important, we expanded the region by adding a 500 bp buffer to each region. These expanded regions of genome were then matched with known gene positions from the Genome Browser, allowing us to associate certain regions with specific genes. Although many of the important regions were not linked to any known genes, we identified plenty known genes for each chromosome and their position in genome using the Genome Browser’s web-API [Kent et al., 2002]. Because the formation of clusters is influenced by important regions of genome, understanding their distribution and relationships can reveal significant biological patterns (e.g., functional similarities) among the samples in the same clusters.

As the hierarchical clustering dendrogram becomes visually intractable for large datasets, we utilized the ETE toolkit to construct a circular dendrogram, which allows for better visual exploration of patterns. In this dendrogram, we highlighted branches in different colors – red, green, blue, and black – depending on the number of important known genes shared by the cluster members. Additionally, we labeled each cluster with pie charts to indicate the number of distinct epigenetic modifiers present in the clusters where more than 30 important genes were found.

We used UMAP to quickly examine how our data clustered, observing whether samples grouped by activity, factor, modifier, or cell type. UMAP’s capability to handle non-metric distances, such as correlation, proved particularly useful in our analysis especially when t-SNE and other non-linear dimensionality reduction techniques usually require metric distances as input.

We set a moderate value of the *min dist* parameter at 0.6 — this parameter controls how tightly UMAP is allowed to pack points together — and a high number of neighbors, 100, to focus on the global structures within the dataset. UMAP’s ability for capturing global structures of dataset effectively allows it to reveal larger-scale patterns, that might not be as clearly visible in methods emphasizing local structures or individual linkages, such as hierarchical clustering.

### E. Co-occurrence analysis of epigenetic modifiers

To analyze data more efficiently, we cut the dendrogram at a specific height so that the correlation distance remains below the threshold of 0.3, ensuring that clusters formed at or below this level had a correlation coefficient of at least 0.7 among their members. We then examined the co-occurrences of epigenetic modifiers in the clusters across all chromosomes. We set a minimum cluster size criterion, considering only those clusters with four or more members. Within these clusters, we determined the frequency at which pairs of unique epigenetic modifiers appeared together. This was accomplished by summing the minimum count of the two modifiers each time they were found within the same cluster. We then plotted these sums for the 50 top co-occurring pairs to visualize the most frequent interactions among the epigenetic modifiers.

### F. Linking with Gene Ontology for functional insights

Building on our analysis, we integrated our findings with external databases to deepen our understanding of the important genes identified across all chromosomes. We used Gene Ontology (GO) and the Kyoto Encyclopedia of Genes and Genomes (KEGG) pathway database for this purpose. KEGG offers a vast collection of databases that shed light on biological pathways and diseases, linking genomic information with higher-order functional insights. In contrast, Gene Ontology provides a well-structured framework for biological activity, defining key concepts used to describe gene function and the relationships among these concepts. It categorizes gene functions into molecular functions, cellular components, and biological processes. The integration with GO and KEGG helped us find out more about what roles these genes might play and how they’re involved in different biological processes and pathways. We used gProfiler tool [Reimand et al., 2007] and Cytoscape [Shannon et al., 2003] to visualize GO and KEGG pathway results, respectively.

## V. Discussion

Hierarchical clustering helped to reveal trends in the epigenome. On a large scale, clusterings from different chromosomes exhibited similar branching statistics. Also, regardless of chromosome, clustering first separated samples into groups based on their epigenetic modifier, and then, at smaller scales, into groups of similar cell types; however, the variation between epigenetic modifiers was larger than the variation between cell types. We also analyzed the adjusted rand index between clusterings on different chromosomes to investigate whether these clusters were similar. The results showed that indeed, there is broad similarity between the epigenetic information on different chromosomes.

We found important regions of the genome in most of the clusters. Through the use of Gene Ontology information, many of the known genes were associated with microRNAs and snoRNAs, which are key to regulating gene expression and thought to be dysregulated during aberrant immune function and in various cancers. Gene ontology revealed several common terms for the important genes found across each chromosome (Fig. 9, 10). These genes are related to chromatin remodeling and nucleosome occupancy, mRNA splicing and regulation, and immune responses. This finding suggests that in addition to the traditional perspective that epigenetic modifiers help regulate the expression of genes involved with cellular processes, they also function in a feedforward manner to regulate their own expression. That is, epigenetic modifications, regardless of a permissive or repressive function, regulate chromatin states for all genes including epigenetic genes. Further, our analysis discovered that processes impacting mRNA regulation, including splicing and microRNAs, are also regulated systematically by epigenetic modifiers. Overall, this suggests that the genes consistently regulated by epigenetic modifiers, regardless of the type of modifier and the extent of permissiveness, are those involved in the formation of chromatin, gene expression processes, and small noncoding RNAs, including microRNAs and snoRNAs. As such, it is possible that identifying epigenetic regulation of noncoding RNAs may be key to understanding disease states, which requires support via empirical data.

The co-occurrence patterns of modifiers showed that Histone 2 and 3 acetylation modifiers commonly co-occurred. The trio of RAD21, SMC3, and CTCF was also frequently found to co-occur. As components of the cohesin complex, RAD21 and SMC3, in conjunction with CTCF, are instrumental in promoting chromatin looping. Several studies have highlighted a correlation between CTCF and cohesin with both the frequency of interaction and gene expression during differentiation, suggesting their significant influence in mediating how chromatin structure affects gene regulation [Phillips-Cremins et al., 2013, Zuin et al., 2014]. The co-occurrence of GTF2F1, TAF1, TBP, POLR2A, and POLR2AphosphoS5 in clusters can be attributed to their involvement in RNA Pol II transcription [Wang et al., 2012, Grau et al., 2015]. EZH2, as well as its phosphorylated form EZH2phosphoT487, were clustered together with SUZ12 and H3K27me3. This aligns with the function of the PRC2 complex in laying down the H3K27me3 mark [Hansen et al., 2008].

Overall, the biologically plausible reasons for the observed clustering of epigenetic modifiers include their involvement in common pathways or biological processes, functional interactions where certain modifiers work together, and their co-binding to similar epigenomic regions. Additionally, post-translational modifications like phosphorylation can link modifiers closely in function and regulation, contributing to their clustering.

In summary, a number of trends were identified by examining whole-genome epigenetic sequencing data for immune cells from the Encode consortium. We observed similar patterns when validating these trends with samples from a different genome assembly, confirming their consistency across assemblies.

We elaborate on the rationale behind analyzing baseline healthy data as a preparatory step for future infectious disease studies.

This analysis serves as a benchmark for understanding the usual clustering patterns of epigenomic features in a non-diseased state. By establishing what constitutes a normal or typical epigenomic configuration, we set a foundation against which changes in the disease state can be measured. This study highlights that certain epigenetic modifiers tend to form clusters in healthy individuals due to their roles in normal physiological processes. These patterns, however, may be disrupted in the context of an infectious disease, leading to altered clustering behaviors. Such changes are indicative not only of disease-specific epigenomic alterations but also of broader biological responses to infection. For instance, the modulation of genes tied to the immune response during an infection could shift the clustering of related epigenetic modifiers away from what is observed in the healthy state. Similarly, epigenetic modifiers that are typically active in maintaining health may exhibit altered behavior under disease conditions. The entropy plots between the healthy baseline and infectious datasets can be examined for variations in entropy levels, along with observed shifts in the overall variability of the dataset.

## VI. Code availability

The code supporting the results of this article is available in the https://github.com/lanl/epigen repository.

## VII. Competing interests

No competing interest is declared.

### A. Funding

The study was supported by the Defense Threat Reduction Agency, Grant DTRA1308139949.

### B. Author’s Contributions

A.K.: formal analysis, data curation, code development, methodology, visualization, writing—original draft. N.L.: conceptualization, formal analysis, methodology, validation, writing—original draft. C.S.: conceptualization, data curation, validation, writing—original draft. K.S.: conceptualization, funding acquisition, supervision, writing—review and editing.

## VIII. Acknowledgements

We thank the LANL Information Science and Technology Institute for providing computational resources.

## References

Tanya Barrett, Tugba O Suzek, Dennis B Troup, Stephen E Wilhite, Wing-Chi Ngau, Pierre Ledoux, Dmitry Rudnev, Alex E Lash, Wataru Fujibuchi, and Ron Edgar. Ncbi geo: mining millions of expression profiles—database and tools. Nucleic acids research, 33(suppl 1):D562–D566, 2005.

Pauline A Callinan and Andrew P Feinberg. The emerging science of epigenomics. Human molecular genetics, 15(uppl 1): R95–R101, 2006.

Nyasha Chambwe, Matthias Kormaksson, Huimin Geng, Subhajyoti De, Franziska Michor, Nathalie A Johnson, Ryan D Morin, David W Scott, Lucy A Godley, Randy D Gascoyne, et al. Variability in dna methylation defines novel epigenetic subgroups of dlbcl associated with different clinical outcomes. Blood, The Journal of the American Society of Hematology, 123(11): 1699–1708, 2014.

Stephen J Clark, Heather J Lee, Sébastien A Smallwood, Gavin Kelsey, and Wolf Reik. Single-cell epigenomics: powerful new methods for understanding gene regulation and cell identity. Genome biology, 17:1–10, 2016.

ENCODE Project Consortium et al. An integrated encyclopedia of dna elements in the human genome. Nature, 489(7414):57, 2012.

Juan Luis Fernández-Morera, Vincenzo Calvanese, Sandra Rodríguez-Rodero, E Menéndez-Torre, and MF Fraga. Epigenetic regulation of the immune system in health and disease. Tissue Antigens, 76(6):431–439, 2010.

Jan Grau, Ivo Grosse, Stefan Posch, and Jens Keilwagen. Motif clustering with implications for transcription factor interactions. Technical report, PeerJ PrePrints, 2015.

Klaus H Hansen, Adrian P Bracken, Diego Pasini, Nikolaj Dietrich, Simmi S Gehani, Astrid Monrad, Juri Rappsilber, Mads Lerdrup, and Kristian Helin. A model for transmission of the h3k27me3 epigenetic mark. Nature cell biology, 10(11): 1291–1300, 2008.

Lifang Hou, Xiao Zhang, Dong Wang, and Andrea Baccarelli. Environmental chemical exposures and human epigenetics. International journal of epidemiology, 41(1):79–105, 2012.

Suganya Ilango, Biswaranjan Paital, Priyanka Jayachandran, Palghat Raghunathan Padma, and Ramalingam Nirmaladevi. Epigenetic alterations in cancer. Frontiers in Bioscience-Landmark, 25(6):1058–1109, 2020.

Peter C Janson and Ola Winqvist. Epigenetics–the key to understand immune responses in health and disease. American Journal of Reproductive Immunology, 66:72–74, 2011.

Randy L Jirtle and Michael K Skinner. Environmental epigenomics and disease susceptibility. Nature reviews genetics, 8(4): 253–262, 2007.

W James Kent, Charles W Sugnet, Terrence S Furey, Krishna M Roskin, Tom H Pringle, Alan M Zahler, and David Haussler. The human genome browser at ucsc. Genome research, 12(6):996–1006, 2002.

Eric B Keverne, Donald W Pfaff, and Inna Tabansky. Epigenetic changes in the developing brain: Effects on behavior. Proceedings of the National Academy of Sciences, 112(22):6789–6795, 2015.

Rasko Leinonen, Hideaki Sugawara, Martin Shumway, and International Nucleotide Sequence Database Collaboration. The sequence read archive. Nucleic acids research, 39(suppl 1):D19–D21, 2010.

I-Hsuan Lin, Dow-Tien Chen, Yi-Feng Chang, Yu-Ling Lee, Chia-Hsin Su, Ching Cheng, Yi-Chien Tsai, Swee-Chuan Ng, Hsiao-Tan Chen, Mei-Chen Lee, et al. Hierarchical clustering of breast cancer methylomes revealed differentially methylated and expressed breast cancer genes. PloS one, 10(2):e0118453, 2015.

Fionn Murtagh and Pedro Contreras. Algorithms for hierarchical clustering: an overview. Wiley Interdisciplinary Reviews: Data Mining and Knowledge Discovery, 2(1):86–97, 2012.

Fionn Murtagh and Pedro Contreras. Algorithms for hierarchical clustering: an overview, ii. Wiley Interdisciplinary Reviews: Data Mining and Knowledge Discovery, 7(6):e1219, 2017.

Yuuki Obata, Yukihiro Furusawa, and Koji Hase. Epigenetic modifications of the immune system in health and disease. Immunology and cell biology, 93(3):226–232, 2015.

Bambarendage PU Perera, Christopher Faulk, Laurie K Svoboda, Jaclyn M Goodrich, and Dana C Dolinoy. The role of environmental exposures and the epigenome in health and disease. Environmental and Molecular Mutagenesis, 61(1):176–192, 2020.

Jennifer E Phillips-Cremins, Michael EG Sauria, Amartya Sanyal, Tatiana I Gerasimova, Bryan R Lajoie, Joshua SK Bell, Chin-Tong Ong, Tracy A Hookway, Changying Guo, Yuhua Sun, et al. Architectural protein subclasses shape 3d organization of genomes during lineage commitment. Cell, 153(6):1281–1295, 2013.

Jüri Reimand, Meelis Kull, Hedi Peterson, Jaanus Hansen, and Jaak Vilo. g: Profiler—a web-based toolset for functional profiling of gene lists from large-scale experiments. Nucleic acids research, 35(suppl 2):W193–W200, 2007.

David B Seligson, Steve Horvath, Tao Shi, Hong Yu, Sheila Tze, Michael Grunstein, and Siavash K Kurdistani. Global histone modification patterns predict risk of prostate cancer recurrence. Nature, 435(7046):1262–1266, 2005.

Paul Shannon, Andrew Markiel, Owen Ozier, Nitin S Baliga, Jonathan T Wang, Daniel Ramage, Nada Amin, Benno Schwikowski, and Trey Ideker. Cytoscape: a software environment for integrated models of biomolecular interaction networks. Genome research, 13(11):2498–2504, 2003.

Takashi Shiina, Kazuyoshi Hosomichi, Hidetoshi Inoko, and Jerzy K Kulski. The hla genomic loci map: expression, interaction, diversity and disease. Journal of human genetics, 54(1):15–39, 2009.

Céline Tiffon. The impact of nutrition and environmental epigenetics on human health and disease. International journal of molecular sciences, 19(11):3425, 2018.

Alexander J Titus, Carly A Bobak, and Brock C Christensen. A new dimension of breast cancer epigenetics. In 9th International Conference on Bioinformatics Models, Methods and Algorithms, 2018.

Arvind K Virmani, Jeffrey A Tsou, Kimberly D Siegmund, Linda YC Shen, Tiffany I Long, Peter W Laird, Adi F Gazdar, and Ite A Laird-Offringa. Hierarchical clustering of lung cancer cell lines using dna methylation markers. Cancer Epidemiology Biomarkers & Prevention, 11(3):291–297, 2002.

Jie Wang, Jiali Zhuang, Sowmya Iyer, XinYing Lin, Troy W Whitfield, Melissa C Greven, Brian G Pierce, Xianjun Dong, Anshul Kundaje, Yong Cheng, et al. Sequence features and chromatin structure around the genomic regions bound by 119 human transcription factors. Genome research, 22(9):1798–1812, 2012.

Alan P Wolffe. Chromatin remodeling: why it is important in cancer. Oncogene, 20(24):2988–2990, 2001.

Yuanxin Xi, Jiejun Shi, Wenqian Li, Kaori Tanaka, Kendra L Allton, Dana Richardson, Jing Li, Hector L Franco, Anusha Nagari, Venkat S Malladi, et al. Histone modification profiling in breast cancer cell lines highlights commonalities and differences among subtypes. BMC genomics, 19(1):1–11, 2018.

Mahdi Zamanighomi, Zhixiang Lin, Timothy Daley, Xi Chen, Zhana Duren, Alicia Schep, William J Greenleaf, and Wing Hung Wong. Unsupervised clustering and epigenetic classification of single cells. Nature communications, 9(1):2410, 2018.

Jessica Zuin, Jesse R Dixon, Michael IJA van der Reijden, Zhen Ye, Petros Kolovos, Rutger WW Brouwer, Mariëtte PC van de Corput, Harmen JG van de Werken, Tobias A Knoch, Wilfred FJ van IJcken, et al. Cohesin and ctcf differentially affect chromatin architecture and gene expression in human cells. Proceedings of the National Academy of Sciences, 111 (3):996–1001, 2014.

